# Is the replication crisis a problem for biologists? A geometric morphometric approach

**DOI:** 10.1101/862052

**Authors:** Juan Vrdoljak, Kevin Imanol Sanchez, Roberto Arreola-Ramos, Emilce Guadalupe Diaz Huesa, Alejandro Villagra, Luciano Javier Avila, Mariana Morando

## Abstract

Replicability of findings is the key factor of scientific reliability. However, literature on this topic is scarce and apparently taboo for large scientific areas. Some authors named the failure to reproduce scientific findings ‘replication crisis’. Geometric morphometrics, a vastly used technique, is especially silent on replication crisis concern. Nevertheless, some works pointed out that sharing morphogeometric information is not a trivial fact, but need to be careful and meticulous. Here, we investigated the replicability of geometric morphometrics protocols on complex shapes and measurement error extension in three different types of taxa, as well as the potentiality of these protocols to discriminate among closely related species. We found a wide range of replication error that contributed from 19.5% to 60% of the total variation. Although, measurement error decreased with the complexity of the quantified shape, it often maintained high values. All protocols were able to discriminate between species, but more morphogeometric information does not imply better performance. We present evidence of replication crisis in life sciences and highlight the need to explore in deep different sources of variation that could lead to low replicability findings. Lastly, we enunciate some recommendations in order to improve the replicability and reliability of scientific findings.

## Introduction

The so-called replication crisis is a hot topic in specialized journals of statistics and psychology [1, 2] and a new field to explore for biologists [3]. The meaning of ‘replication crisis’, in broad sense, is associated with the failure to reproduce results of studies. However, most scientific researches never attempt to replicate results, possibly because – fed by the ‘publish or perish’ dogma – most scientific journals have within their scopes explicit policies against publishing replication studies [4]. Non-replicability leads to lack of reliability in scientific findings because it compromises our belief on the generality of scientific theories.

Publication bias, questionable research practice (QRP) and over-confidence on null hypothesis significance test (NHST) are bad practices that affect replicability without threatening the generality of scientific facts [2, 5]. In addition to the rejection of replication articles, the strong tendency to publish only significant results is the second factor that influences publication bias [6, 7, 8]. QRP refers to a set of post-hoc decisions that include: data point exclusion to improve statistical significance, stopping data collection because results show significant differences, no report of parameters that were statistically non-significant, among others [3].

The NHST and the p-value thresholds are the current paradigms for research, publication and discovery in biological and social sciences [9, 10]. This set of ideas leads to several mistakes and could be the cause of publication bias and QRPs. Among the main mistakes we can mention: the dichotomization of results into “significant” and “no significant”; focus only on significant results even when they are irrelevant (e.g. descriptive statistics); ignore other evidence such as magnitude of effect; several misinterpretations of p-value; and the implausibility of null hypothesis when the effects are small, because the possibilities of systematic bias and variation due to highly variable measurements could result in similar small effects [11, 12].

Measurement error (ME) is an uncontrolled variation that could aggravate the replication crisis [1]. Given its random nature, ME is frequently associated with noise around the true values. Thus, if an effect is found in a noisy statistical environmental, then it is logical to think that the actual effect is really strong [13]. However, effect size estimation can be exaggerated and the outcomes can result biased by a poor measurement [1, 14].

Geometric morphometric is a simple technic to quantify, identify and describe shapes independently of size. Thus, three steps are necessary to obtain morphometric data: photograph the object, placement of landmarks (or outliner contours) in anatomical positions and superimposition of these points [15, 16]. There are a few dozens of articles that help geometric morphometric operators to guide and improve their analysis [17, 18, 19, 20] and at least 21,500 articles that made use of this technique (according to a brief search in Academic Google). However, little is known about the source of variation that could generate spurious results [21, 22, 23, 24].

In this sense, Fruciano (2016) reviewed the common sources of error in geometric morphometrics with emphasis on ME. He enunciated different forms to assess ME and concluded that researchers have to take into account certain considerations that compromise accurate measurement, e.g. effort invested in digitalization of images [25], trade-off between sample size and specimen quality [21, 26], maintenance of coplanarity in 3D structure [27], among others. In complex shapes, several conflictive points could lead to overinflate ME due to low accuracy landmarks [28] or high landmarking bias [29]. Moreover, a good treatment of conflictive points could be the cornerstone to increase replicability in geometric morphometrics [30].

The term complex shapes refer to certain configurations where the placement of landmarks is not trivial. In this regard, Bookstein (1991) described type I, II and III landmarks according to a scale from more to less clarity of the anatomical point, respectively [31]. Several authors reported that type III landmarks are clearly associated with high ME [29, 32, 33, 34]. Given that the analytical procedure is the same for these types of landmarks and that this distinction has a strong subjective or arbitrary character (35), several articles do not make use of this distinction. On the other hand, following the aim to describe complex shapes, curves or contour are more suitable because point-to-point homology is hard to ensure [36, 37]. However, no measurement technique is error free. In fact, there is a positive relationship between the number of semilandmarks used to describe a curve and the ME [38]. Therefore, there is a trade-off between the ME and the potential of description in these techniques.

Here, we carry out the first study to analyze the extent of replication crisis in life sciences using geometric morphometrics, which is a widely used tool in biology and anthropology. Different spurious (later called extrinsic) sources of variation were analyzed. In this sense, the principal aim of this work was to quantify these sources of variation through different geometric morphometric methods in lizards and discuss the principal implications for biological inferences. Additionally, we evaluated how ME extend to other taxa (a fly and a plant) and assess the potentiality of each geometric morphometric method to describe and discriminate among closely related species. Finally, we advocate the use of a clear and solid statistic framework without falling into the apparent need of QRP or NHST in order to respond these aims.

## Materials and methods

### Experimental designs, sample, data collection and methodological approach

The following three designs were developed in order to address three different objectives: The first objective was to analyze different factors that may affect replication. To address this, 25 photographs of male lizards *Liolaemus elongatus* were mirrored and then landmarked/outlined each side twice by five operators: among-operators design. The second objective was to estimate ME across three very different taxa: *L. elongatus* lizards, the fly *Drosophila buzzatii* [39] and the grape *Vitis riparia* (data available on https://dataverse.harvard.edu/dataverse/VitisLeafVariation, [40]). In this case, each of 25 photographs was landmarked/outlined in quadruplicate and analyzed each taxon separately: across-taxa design. The third objective was to assess the potentiality of each morphometric configuration (described below) to differentiate among closely similar shapes. Thus, 25 male specimens of three closely related lizard species (*L. elongatus*, *L. shitan* and *L. choique*; [41]) with several morphological similarities were landmarked/outlined: related-species design (details of specimen’s voucher numbers, collection locality and other data on appendix 1).

We took dorsal photographs of the head of each specimen using a Canon 1000D camera mounted in a fixed tripod. For flies, we removed left wing, mounted them on slides with DPX and photographed them at 40x magnification using a digital camera attached to a microscope (Nikon E200). To characterize the shape, we placed landmarks in four different configurations using TpsDIG 2.31 [42]. Shape variation was estimated by general Procrustes analysis [43, 44], and then we performed principal component analysis to summarize the information of shape in uncorrelated form. In addition, we employed another approach to quantify shape variation based on elliptic Fourier descriptors [45]. In this sense, outlines from digital images were used to obtain Fourier coefficients normalized for size, rotation and starting point, then we built a variance-covariance matrix that was used as input in a principal components analysis. Morphometric analyses were performed with R statistical software [46] packages Momocs [47] and geomorph [48].

For each of the three designs, we developed five morphometric protocols: two landmark-only protocols with six and ten landmarks for lizards and leaves and ten and fifteen landmarks for flies (P-L and F-L protocols for partial-landmark and full-landmark respectively); two semilandmark protocols [37], both starting from the same P-L configuration, with one and two curves (P-S and F-S protocols for partial-semilandmark and full-semilandmark respectively). In both cases, partial protocols (P-L and P-S) were less time expensive and could explain more stable and homologous onfiguration but less explanatory power than full protocols (F-L and F-S). It should be noted that the partial methods are a subset of full methods. Lastly, we used a contour protocol, a novelty in herpetological research.

We examined one-side morphologies except in the cases of lizards’ contours in the across-taxa and related-species designs, where the whole pineal scale was used. In the case of lizards’ contour in the among-operators design, half of pineal scale and whole parietal scale were outlined (Appendix 2 includes details of contours and landmark configuration; also see [40, 49, 50]).

To carry out the among-operators design, the order of the five protocols was randomly selected for each operator. Also, in order to represent the greatest possible variation in ME, operators with different degree of knowledge on morphometric techniques were chosen. For the other two designs (across-taxa and related-species), only one of the operators performed all five protocols. Finally, we computed the data gathering processing time for each protocol applied to ten specimens randomly selected from each taxon.

### Models and statistical analysis

Among-operators design: we used hierarchical models to estimate seven variance effects: specimen variation, operator variation, the interaction with side and between them, resulting in specimen*operator, specimen*side, operator*side and specimen*operator*side variation and, finally, ME. Specimen and specimen*side variation (latter known as fluctuation asymmetry) are two intrinsically natural sources of variation (intrinsic variation), while the other variance effects depend on operator errors, biased measurement, and consistency of these (extrinsic variation, composed of ME and replication factors). As a result of this model, we evaluated different factors affecting the replication (operator or operator interaction effects), and thus reliability of the measurement technique in a relative manner with intrinsic variation. Across-taxa design: to evaluate ME of each of the five protocols among taxa as accurately as possible, we used hierarchical models that included only specimen variance and ME, i.e. each taxa was analyzed separately. In this sense, each protocol was applied to grape leaves, fly wings and lizard heads by one operator. Related-species design: we analyzed the effects of morphological differences among species (details of models on appendix 3).

It is simple to predict that the placement of more points implies more processing time. For this reason, we employed a linear regression between processing time and centroid size. Centroid size is more suitable to explain processing time than the simple sum of landmark and semilandmark points, since it is a good proxy of number of points by definition (the square root of the sum of the squared distances of each landmark to the centroid configuration, [16]), and the operators could spend more time in mouse displacement in large than small sizes with equal number of points.

In the described protocols, except for the related-species design where we explored the minimal number of principal component (PC) axes needed to differentiate among species, we employed the first PC axes that explain at least 60% of the total variation to perform statistical analyzes. Given the large number of PC axes (8 to 48), our decision criterion was taken to explain more than half of the total variation. Furthermore, we also investigated the rest of the PC axes in search of substantial morphological changes to incorporate into statistical analyzes; however, we did not find such changes in those PC axes.

The models were fitted within a Bayesian framework that eased implementation of variance components and its uncertainty. Posterior distribution of parameters were estimated using three independent Markov Chain Monte Carlo (MCMC) runs for 100,000 iterations and 20% of burn-in each implemented in JAGS 4.3.0 [51] using the R packages jagsUI [52] and rjags [53]. The observations were centered and standardized to reduce autocorrelation of chains [54]. Convergence was assessed using Gelman and Rubin statistics Ȓ [55] and by visual inspection of trace plots. We used weakly informative prior distributions to include small amounts of information on parameter and hyperparameter such as non-negative possibilities and to avoid meaningless values [56, 57]. Finally, we denoted differences between two samples of the response variable as standardized difference, whereas we reserved effect size to the distribution of the standardized differences from posterior distribution ([58] and analytical according to [59]) and reported mean and High Posterior Density interval (HPD) using the R package coda [60].

## Results

### Among-operators design

We found extrinsic variation with all five protocols considering the first PC that explains the greatest morphological variation (Fig 1). While the contour protocol had the best performance (highest sources of intrinsic variation and smallest sources of extrinsic variation), the other protocols showed a trade-off among different sources of variation. In this sense, both partial protocols showed the highest levels of variation among operators and explained very similar intrinsic variation with a greater uncertainty in P-S protocol. Full protocols explained more intrinsic variation than the previous, where F-S protocol captured more extrinsic variation than F-L protocol. As a good proxy of measurement bias, we found high levels of operator*specimen variation in F-L protocol due to the consistent placement of two conflictive landmarks on some specimens (Fig S1.a).

**Figure 1:**
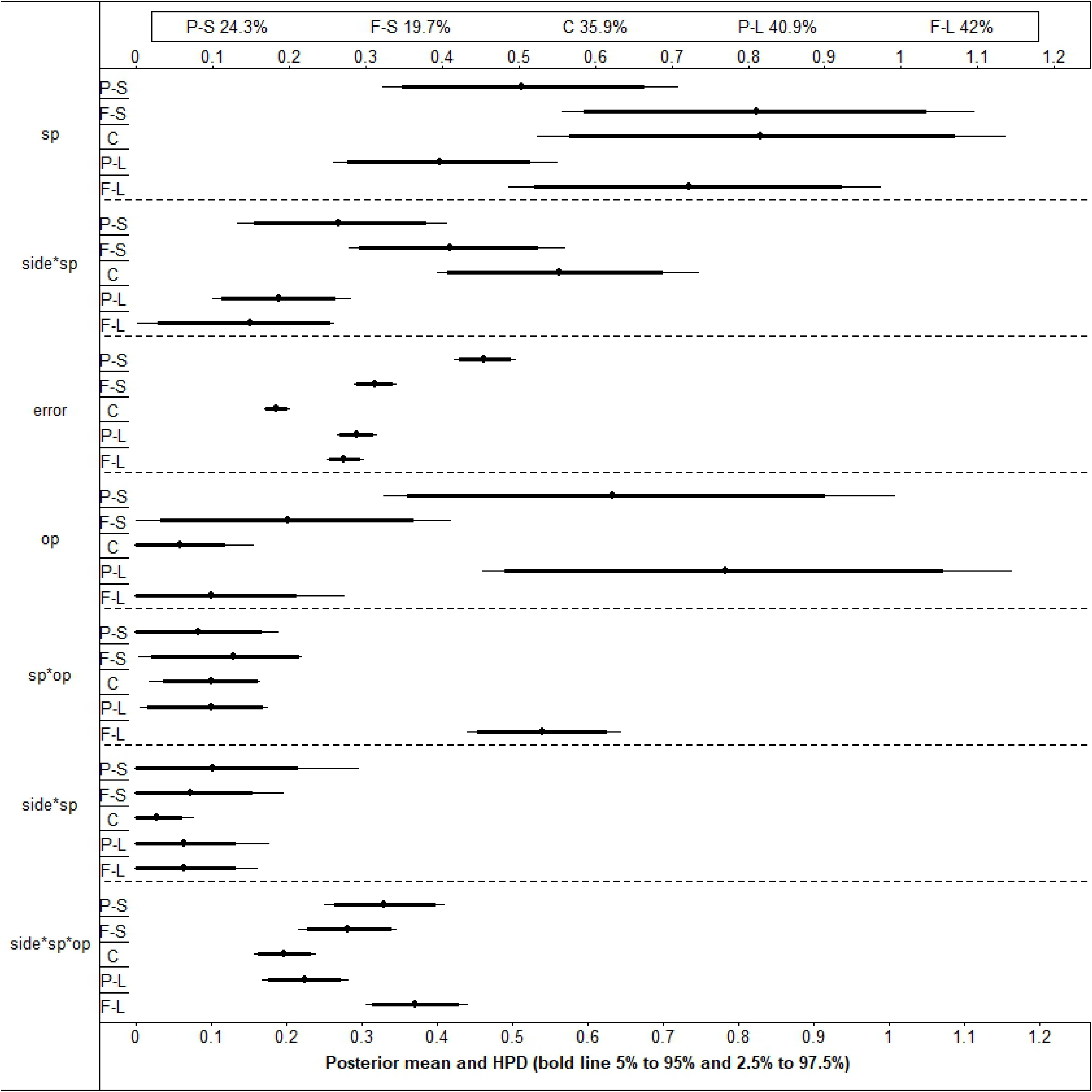
Posterior mean and High Posterior Density interval (HPD) of 90% (bold line) and 95% (thin line) for each source of variation: sp: Specimen, side: Side, op: Operator, error: Measurement error. * denote interaction between sources of variation. Protocols: P-S: Partial-Semilandmark, F-S: Full-Semilandmarks, C: Contour, P-L: Partial-Landmark, F-L: Full-Landmark. Box of the top, percentage of variation explained by the first principal component for each protocol.

Replication error was always greater than ME for all protocols (see PC1 in Fig 2). More than half of the total variation was explained by replication factors in P-L protocol (56.7%), closely followed by F-L and P-S with almost half of the total variation (48% and 47.7%, respectively). In contrast, replication error contributed 30.4% of mean variation to the whole model in F-S and remarkably less in contour protocol (19.6%). Nevertheless, ME explained no negligible variation in all protocols. In this sense, P-S expressed noticeably greater variation of ME (19.5%) than contour protocol (9.5%), whereas P-L, F-L and F-S showed similar variation (14.4%, 12.4% and 14.2%, respectively).

**Figure 2:**
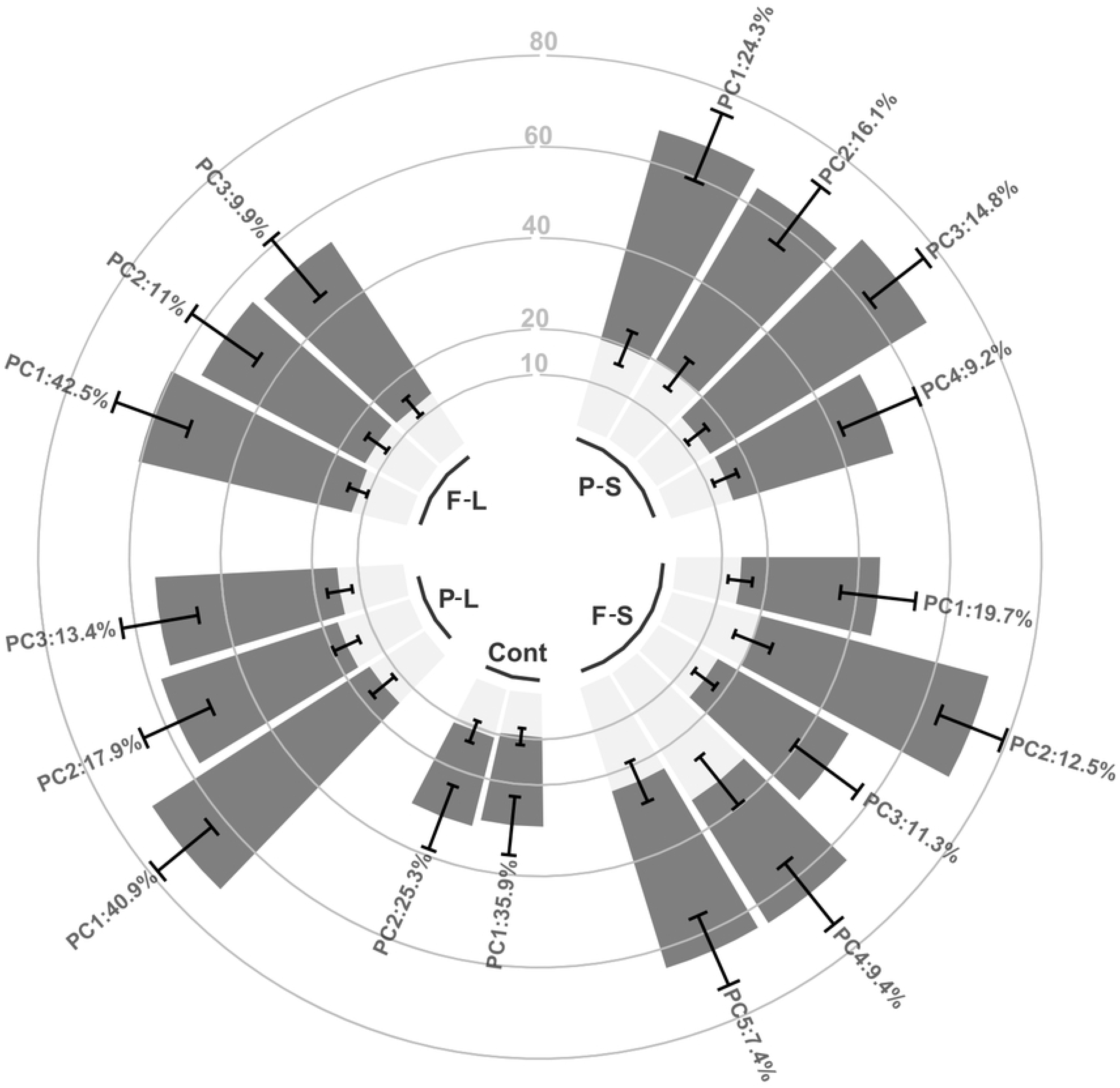
Barplot of posterior mean and High Posterior Density interval (error bars) of 95% measurement error (light gray bars) and replication factors (dark gray bars). Protocols: P-S: Partial-Semilandmark, F-S: Full-Semilandmarks, Cont: Contour, P-L: Partial-Landmark, F-L: Full-Landmark. Percentage of variance explained by each principal component analyzed on top of the bars. ggplot2 (Wickham, 2016) was used to develop this figure.

Given all PCs analyzed, contour protocol maintained lowest mean values of extrinsic variation (Fig 2). More than 60% of the total variation was explained by extrinsic factors in the first three PCs of the P-S protocol. In F-S protocol, the first and third PCs showed smaller extrinsic variation than the others. Both P-L and F-L protocols improved the levels of extrinsic variation through the PCs, however high levels of operator*specimen variation were found in PC2 of the former protocol (Fig S2), due to a consistent bias on some specimens (Fig S1.b).

### Across-taxa design

This design exposed at least three clear patterns (Fig 3). First and more conspicuous, ME variation resulted highest in lizards, followed by flies and finally leaves (the averages of the ME variation weighted by the morphological variation explained by each PC were 28.2%, 9.8% and 2.6%, respectively). In particular, the protocol with greatest ME contributed 57.3%, 19.8% and 7.6% to the total variation while the protocol with smallest ME contributed 15%, 11.3% and 1.5% to the total variation in lizards, flies and leaves respectively. Thereby, we found a wide range of ME dependent on both taxa and protocol, meaning that some protocols are more suitable for one taxon than other.

**Figure 3:**
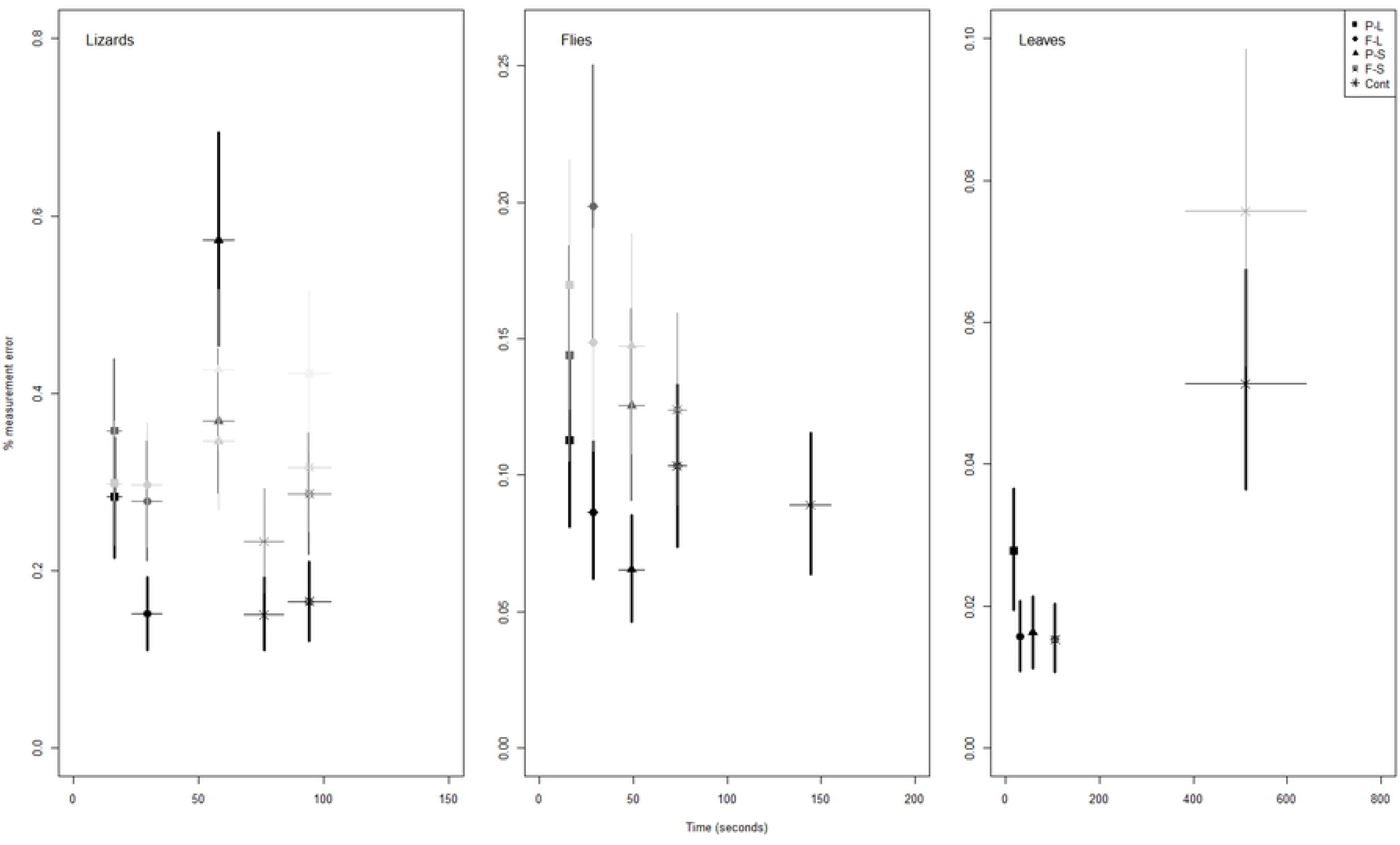
Posterior mean and High Posterior Density interval (error bars) of 95% measurement error vs mean and standard deviation of processing time. Protocols: P-S: Partial-Semilandmark, F-S: Full-Semilandmarks, Cont: Contour, P-L: Partial-Landmark, F-L: Full-Landmark. The Principal Components are represented by a gray shade scale, where the darker points and lines correspond to the first PC and the subsequent ones increasingly clearer.

Second, as we expected, processing time was longest in protocols with more points (understanding points as a number of landmarks plus semilandmarks), i.e. the processing time for all taxa follows from longer to smaller: F-S, P-S, F-L and P-S. Indeed, we found a positive correlation between processing time and specimen size across protocols (excluding contour protocol for the analysis, Fig S3).

Third and more interesting, contour protocol showed an independent pattern of processing time with respect to the other protocols. In this sense, contour processing time resulted smaller than F-S protocol in lizards but higher than all protocols in the other taxa. Moreover, the difference between contour and F-S protocols mean time elapsed resulted in 0.8, 1.96 and 4.87 (but 2.12, 7.89 and 4.29 of standardized differences) for lizards, flies and leaves respectively.

### Related-species design

All protocols were able to discriminate between species more or less clearly (Table 1). In this sense, both semilandmark protocols presented highly clear differences among species with a slightly better performance in the partial protocol. However, the morphological information explained by these protocols resulted redundant (Fig S1c) and more time expensive in the full protocol. The landmark protocols also showed highly clear differences among species, but each protocol explained dissimilar morphological information (Fig S1d). It is critical to point out that main differences between species for PC1 of F-L protocol were due to changes on the same conflictive landmarks that were found that strongly biased the among-operators design (Fig S1a). Contrastingly, the differences among species found by contour protocol were slightly less clear than the previous mentioned, indeed it was necessary to seek in more than 2 PC axes.

**Table 1:**
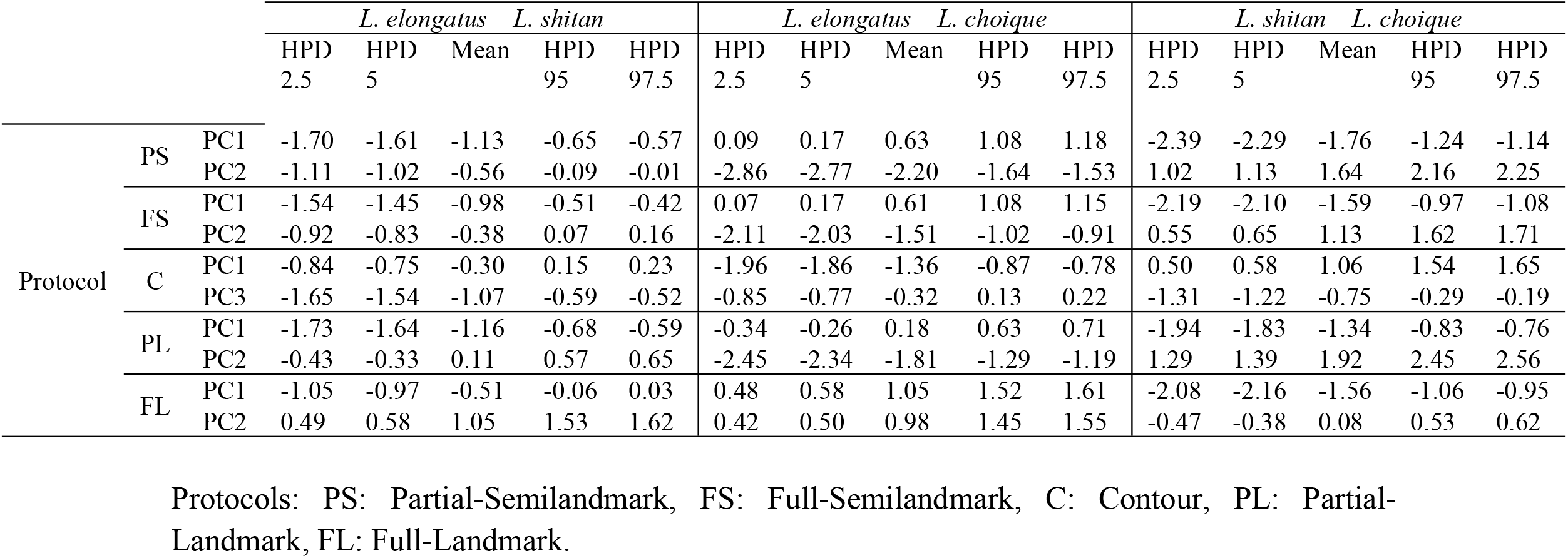
Mean values, High Posterior Density interval (HPD) of 95 and 90% of the effect size distribution resulting from species comparisons. Number of Principal Components (PC) analyzed to reach a clear differentiation between species.

|  |  |  | *L. elongatus – L. shitan* |  |  |  |  | *L. elongatus – L. choique* |  |  |  |  | *L. shitan – L. choique* |  |  |  |  |
| --- | --- | --- | --- | --- | --- | --- | --- | --- | --- | --- | --- | --- | --- | --- | --- | --- | --- |
|  |  |  | HPD<br>2.5 | HPD<br>5 | Mean | HPD<br>95 | HPD<br>97.5 | HPD<br>2.5 | HPD<br>5 | Mean | HPD<br>95 | HPD<br>97.5 | HPD<br>2.5 | HPD<br>5 | Mean | HPD<br>95 | HPD<br>97.5 |
| Protocol | PS | PC1 | -1.70 | -1.61 | -1.13 | -0.65 | -0.57 | 0.09 | 0.17 | 0.63 | 1.08 | 1.18 | -2.39 | -2.29 | -1.76 | -1.24 | -1.14 |
|  |  | PC2 | -1.11 | -1.02 | -0.56 | -0.09 | -0.01 | -2.86 | -2.77 | -2.20 | -1.64 | -1.53 | 1.02 | 1.13 | 1.64 | 2.16 | 2.25 |
|  | FS | PC1 | -1.54 | -1.45 | -0.98 | -0.51 | -0.42 | 0.07 | 0.17 | 0.61 | 1.08 | 1.15 | -2.19 | -2.10 | -1.59 | -0.97 | -1.08 |
|  |  | PC2 | -0.92 | -0.83 | -0.38 | 0.07 | 0.16 | -2.11 | -2.03 | -1.51 | -1.02 | -0.91 | 0.55 | 0.65 | 1.13 | 1.62 | 1.71 |
|  | C | PC1 | -0.84 | -0.75 | -0.30 | 0.15 | 0.23 | -1.96 | -1.86 | -1.36 | -0.87 | -0.78 | 0.50 | 0.58 | 1.06 | 1.54 | 1.65 |
|  |  | PC3 | -1.65 | -1.54 | -1.07 | -0.59 | -0.52 | -0.85 | -0.77 | -0.32 | 0.13 | 0.22 | -1.31 | -1.22 | -0.75 | -0.29 | -0.19 |
|  | PL | PC1 | -1.73 | -1.64 | -1.16 | -0.68 | -0.59 | -0.34 | -0.26 | 0.18 | 0.63 | 0.71 | -1.94 | -1.83 | -1.34 | -0.83 | -0.76 |
|  |  | PC2 | -0.43 | -0.33 | 0.11 | 0.57 | 0.65 | -2.45 | -2.34 | -1.81 | -1.29 | -1.19 | 1.29 | 1.39 | 1.92 | 2.45 | 2.56 |
|  | FL | PC1 | -1.05 | -0.97 | -0.51 | -0.06 | 0.03 | 0.48 | 0.58 | 1.05 | 1.52 | 1.61 | -2.08 | -2.16 | -1.56 | -1.06 | -0.95 |
|  |  | PC2 | 0.49 | 0.58 | 1.05 | 1.53 | 1.62 | 0.42 | 0.50 | 0.98 | 1.45 | 1.55 | -0.47 | -0.38 | 0.08 | 0.53 | 0.62 |

Protocols: PS: Partial-Semilandmark, FS: Full-Semilandmark, C: Contour, PL: Partial-Landmark, FL: Full-Landmark.

## Discussion

We analyzed several factors of replication error and ME in geometric morphometrics, and also the potentiality of each developed protocol. We found worrying levels of extrinsic variation across the whole study, highlighting the need of in depth inquiry on replication crisis in life sciences. Moreover, we have shown that conformational changes with a high risk of measurement bias might exaggerate the true morphological differences, further aggravating concern about replication crisis.

In addition, little is known about the reproducibility of geometric morphometrics results, much less how the decision making impacts on these results. Fagertun et al (2014) [30] reported that the operator variation was associated to particular landmarks (also reported by [28, 61, 62]) and that such variation resulted similar to variation among individuals. However, what they called error term (it was not the ME by model construction) resulted twice than each of the previous mentioned variation, while Robinson and Terhune (2017) [62], Fruciano et al (2017) [63] and Shearer et al (2017) [64] found that highest variation was attributable to inter-operator factor. In agreement with these last three works, we show that the replication error (i.e. inter-operator factors) was always greater than ME, and that in most cases, the total extrinsic variation resulted greater than intrinsic variation. A clear operational conclusion should be digitizing on original images by one operator rather than utilizing data sets developed by more than one operator [63, 64]. Moreover, replication factors accounted for at least 19% of the total variation and rose to almost 60% in the less replicable protocol. In this regard, we also want to point out that the replication crisis is a fact in life sciences [3, 65, 66].

Bias could be defined as systematic error. Unlike any random error, measurement bias could lead to mean differences between groups when this does not exist. However, in geometric morphometrics, Fruciano et al (2017) [63] showed that bias accounts for a small proportion of variation and becomes significant when highly variable landmarks were removed. We found biased measures on two different protocols: F-L and P-L. Variation due to these biased measures were captured by PC1 (42.5%) and PC2 (16.1%), respectively. Curiously, *L. choique* was differentiated from the other species mainly by morphological changes around the two conflictive landmarks involved in the measurement bias of F-L protocol. If the operator’s experience may influence in the degree of biased measurement [64], then bias on F-L protocol could become seriously problematic.

Measurement error is a widely studied issue in the scientific literature and a concern for a large percentage of publications [14]. Some authors predict that with technological advances, ME would probably become a less frequent problem but the large amount of data available obtained by other researchers could incorporate new sources of variation [63, 67, 68]. Our findings indicate that there is a relationship between complex shapes and ME. In this way, photographed lizards had some broken or missing scales and colors that made difficult the digitalization. The width of *Drosophila*’s veins might be the key factor of the ME levels found here, because the intersection of them is not clear. By last, leaves had high specimen variation and clear positions to landmarks or contours.

Another key factor in deciding how to digitize samples is the processing time. Despite the fact that this factor resulted similar among each protocol and taxon, the contour protocol showed a distinctive pattern: the processing time was positively correlated with size and complexity. In this sense, the effect of size is expressed in the differentiation between flies and lizards, where the contour of the former occupied almost the entire image while the contour of the latter occupied a little place in the image. On the other hand, the flies’ wing is an appendix more or less round, and clearly distinguishes itself from the innumerable grape leaf peaks, and in this sense we described the difference in the processing time due to complexity (appendix 2).

Geometric morphometrics is ubiquitous, well accepted and a practical tool to quantify morphological phenotypes [69, 70, 71], fluctuating asymmetry [72, 73], acoustic signals [74], useful forensic patterns [75, 76] among others. Selecting a configuration that faithfully represents the shape analyzed is an obvious but not a trivial notion. Here, we studied the potentiality of each protocol to discriminate among species and found that more landmark points does not necessary explain more shape information. Indeed, P-S resulted better than F-S protocol to discriminate among species (Table 1). F-L protocol also differentiates species with high performance; however its relationship with measurement bias detracts from this differentiation (Fig S1a). Despite contour protocol expressed differences at one scale level, species discrimination was successful highlighting that this method deserves to be studied in depth for its high performance in all designs.

Certain recommendations should be noted. First and foremost, each morphogeometric stage (up to results) must be developed by one person. The great variation found in this work was only the result of placement of landmarks by five operators. If other five operators had photographed each or some specimens, for instance, then the extrinsic variation should be greater. Second, search in bibliography and select homologous positions for landmarks placement are good practices to improve replication. Moreover, pioneer morphometric studies need to be more careful and seek the most stable landmarks configuration by pilot tests. Third, quantify ME and, if possible, add to the whole model. There are many ways to estimate ME in geometric morphometrics [24], but most of them entail an extra effort such as multiple digitizations, learning about novel methods, good data management, among others; instead of this, most researchers prefer to focus their efforts on expanding their dataset. Fourth, select a method that has a high quality-processing time ratio. Sometimes, long processing time can enhance the ME. Fifth, complex forms do not necessarily need complex landmarks conformation. We have shown that there are not many differences between the “resolution” of partial and full protocols, but the latter needs considerable more processing time. Sixth, be careful (or be Bayesian) when the underlying effect is small and sampling error is large, because experiments that achieve statistical significance must have exaggerated effect sizes and are likely to have the wrong sign [77].

Overall, our results call researchers to reflect on their conclusions’ extent and what this implies, for instance, in the widespread discourse of scientific truth and scientific unity [78]. Moreover, this problem could get worse if we take into account some of the current proclamations about the role of subjectivity in the scientist’s tasks, for example the criticism developed by Garnett and Christidis (2017) [79] on the arbitrariness of taxonomy (but see [80, 81, 82]). We invite other researchers to repeat this kind of assay in their disciplines to understand how deep is the crisis of replication in the natural sciences.

## Authors’ contribution

JV designed the experiment, performed the analyses and drafted the manuscript. JV, KIS, RAR, EDH and AV generated the dataset. JV, KIS, and MM corrected and subsequently rewrote the manuscript. LJA and JV contributed to field sampling. All authors gave final approval for publication.

## Funding

Financial support was provided by the following grants: ANPCYT-FONCYT 1397/2011 (LJA); ANPCYT-FONCYT 1252/2015, PIP-CONICET 0336/13 (MM), and a doctoral fellowship (JV) from Consejo Nacional de Investigaciones Científicas y Técnicas (CONICET).

## Acknowledgements

We thank members of the Grupo de Herpetología Patagónica (GHP) for assistance in field collections, in the laboratory, and/or assistance in animal curation procedures. We thank the fauna authorities from Río Negro, Neuquén, Mendoza, and Chubut Provinces for collection permits.

---

**Figure S1:**
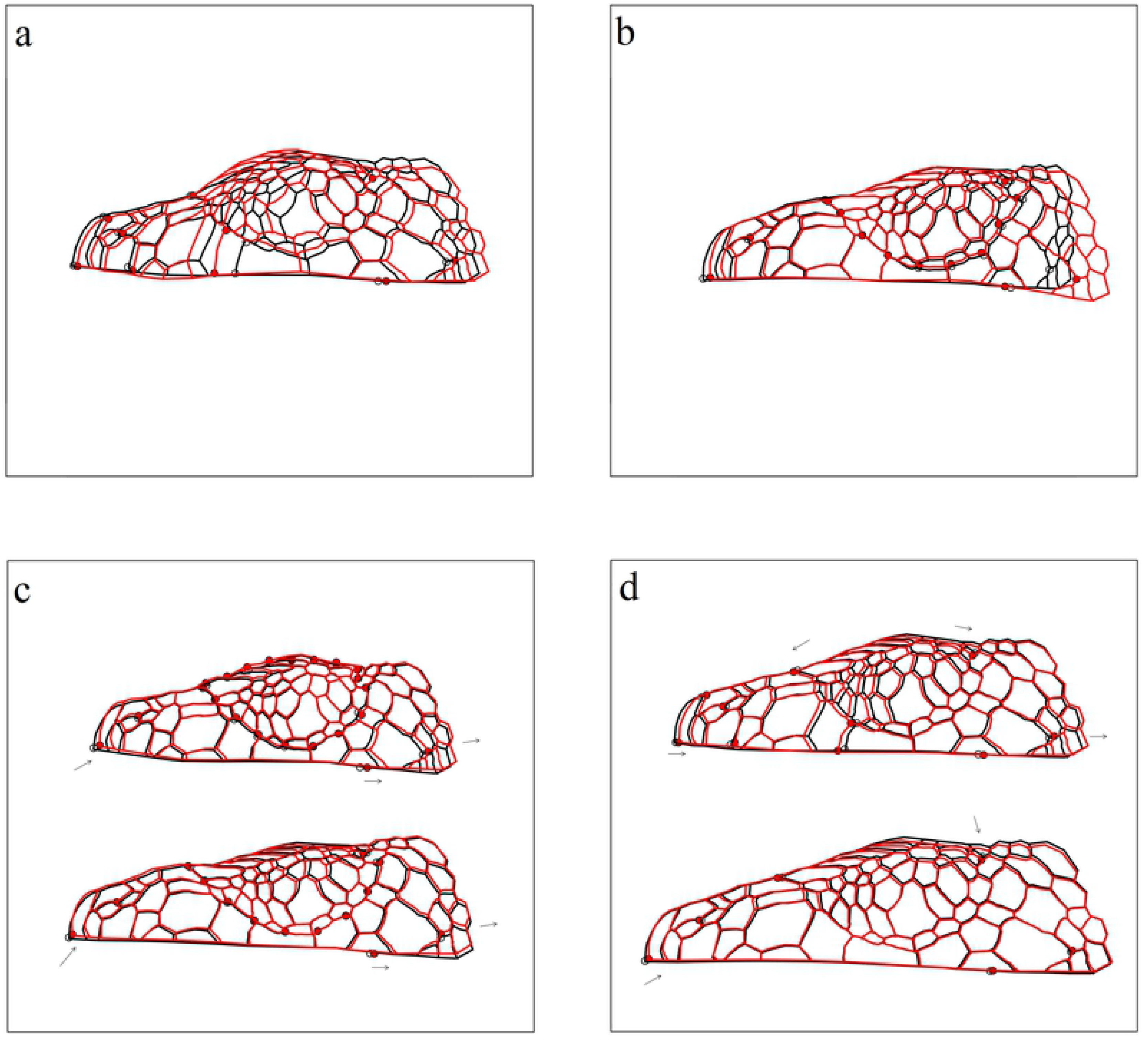
Different shape changes. a) Changes between the same specimen digitized by different operators on Full-Landmark protocol. b) Changes between the same specimen digitized by different operators on Partial-Landmark protocol. c) Changes between consensus and a randomized specimen on Full-Semilandmak (top) and Partial-Semilandmark (down) protocols. Note the change similarities between protocols (marked with arrows). d) Changes between consensus and a randomized specimen (same that c for a better compression) on Full-Landmak (top) and Partial-Landmark (down) protocols. Note the change dissimilarities between protocols (marked with arrows).

**Figure S2:**
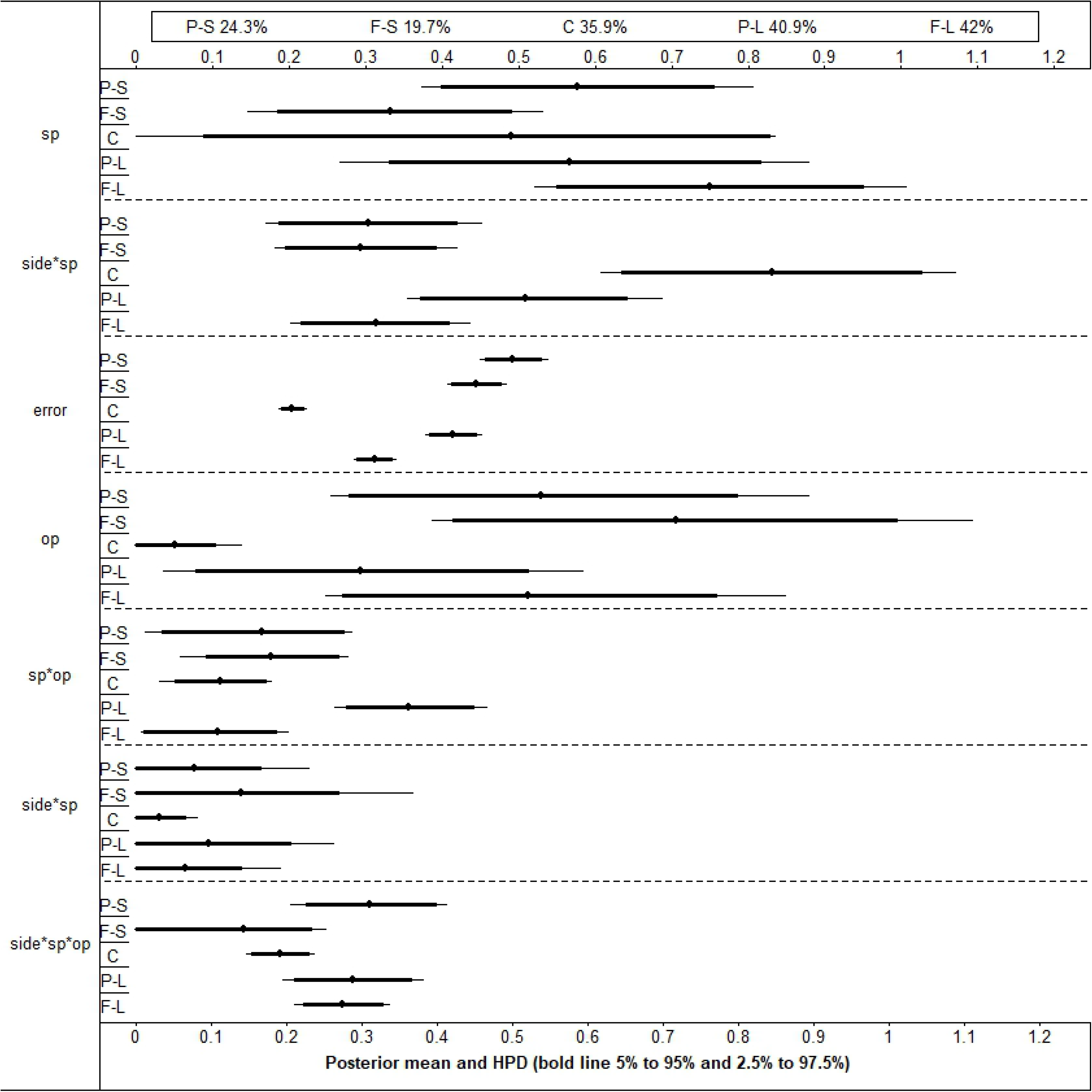
Posterior mean and High Posterior Density interval (HPD) of 90% (bold line) and 95% (thin line) for each source of variation: sp: Specimen, side: Side, op: Operator, error: Measurement error. Aesthetics denote interaction between sources of variation. Protocols: P-S: Partial-Semilandmark, F-S: Full-Semilandmarks, C: Contour, P-L: Partial-Landmark, F-L: Full-Landmark. In the box of the top, percentage of variation explained by the second principal component for each protocol.

**Figure S3:**
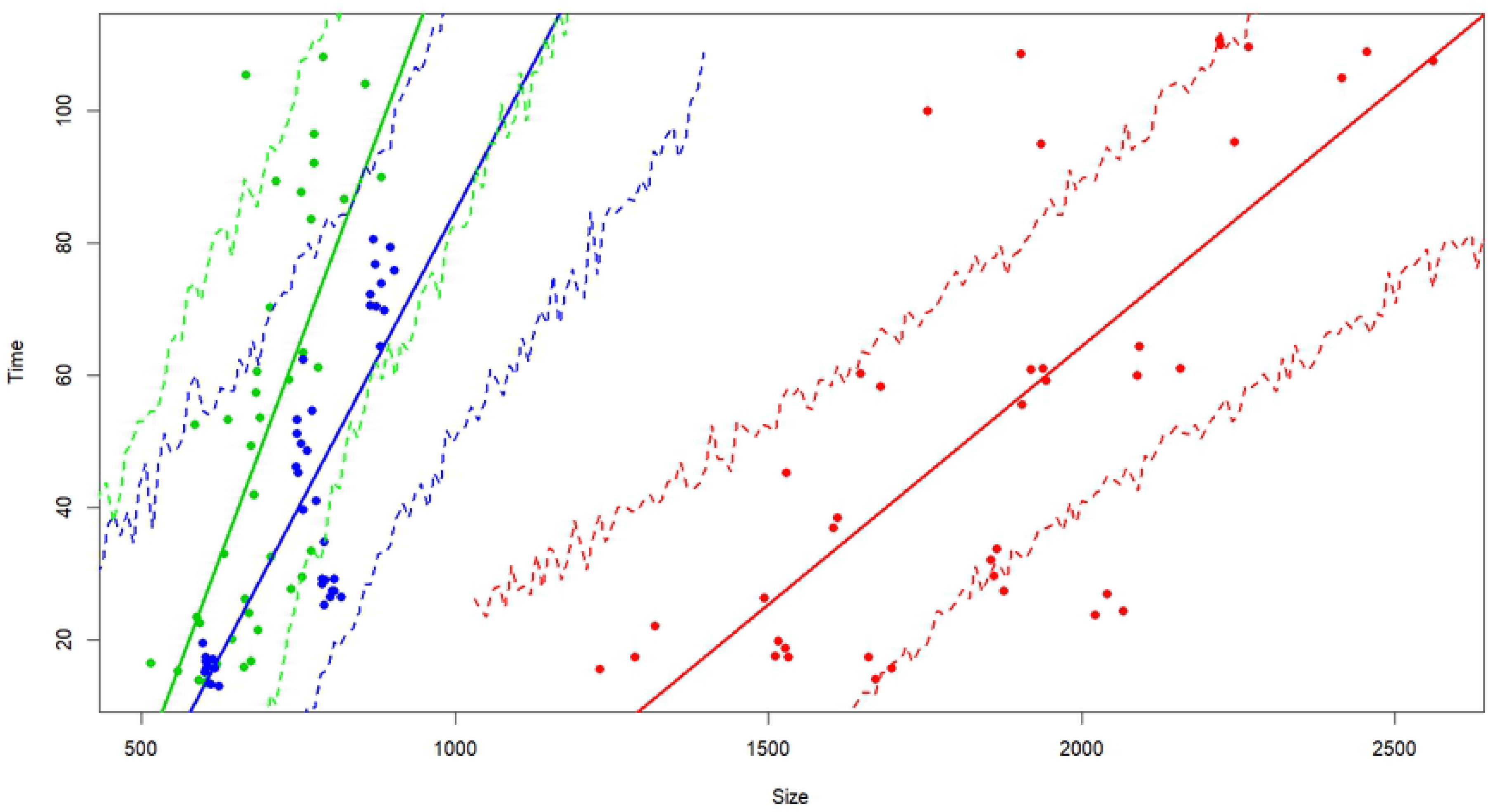
Correlation between time (in seconds) and size for landmark and semilandmark protocols in each taxon. Green: lizards (*Liolaemus elongatus*); blue: flies (*Drosophila buzzatii*); red: leaves (*Vitis riparia*). Continuum lines represent the lineal regression whereas dashed lines represent the simulation of credibility interval.

